# Inflammatory bowel disease microbiotas alter gut CD4 T-cell homeostasis and drive colitis in mice

**DOI:** 10.1101/276774

**Authors:** Graham J. Britton, Eduardo J. Contijoch, Ilaria Mogno, Olivia H. Vennaro, Sean R. Llewellyn, Ruby Ng, Zhihua Li, Arthur Mortha, Miriam Merad, Anuk Das, Dirk Gevers, Dermot P.B. McGovern, Namita Singh, Jonathan Braun, Jonathan P. Jacobs, Jose C. Clemente, Ari Grinspan, Bruce E. Sands, Jean-Frederic Colombel, Marla C. Dubinsky, Jeremiah J. Faith

## Abstract

To examine the functional contribution of Inflammatory Bowel Disease (IBD) microbes to immune homeostasis and colitis, we colonized unchallenged and colitis-susceptible germ-free mice with over twenty human intestinal microbiotas from healthy and IBD donors. Compared to healthy microbiotas, IBD microbiotas led to expanded RORγt^+^Th17 cells and reduced RORγt^+^Treg in the gut of unchallenged gnotobiotic mice and increased disease severity in colitis-susceptible mice. The proportions of RORγt^+^Th17 and RORγt^+^Treg induced by each microbiota were highly predictive of the human disease status and strongly correlated with disease severity in colitis-susceptible mice colonized with the same human microbiotas. The transmittable functional potential of IBD microbes suggests a mechanism for a microbial contribution to IBD pathogenesis and a potential route for its treatment and prevention.

---

Culture-independent studies have revealed dramatic interpersonal variation in the structure and composition of the human gut microbiome [1]. How this interpersonal microbiome variation influences health and susceptibility to disease remains a largely unknown but essential question to translate the results of microbiome research to health benefits. Numerous animal models suggest the microbial strains that constitute the gut microbiota play a crucial role in shaping the immune system, consequentially influencing host susceptibility to autoimmunity, inflammatory disease and infection [2-5]. Inflammatory bowel disease (IBD), including Crohn‘s disease (CD) and ulcerative colitis (UC), is associated with an altered intestinal microbiota [6-8] and genetic defects in microbial handling are implicated in disease [9]. Therefore, interventions targeting the microbiota are a potential path for IBD treatment [10, 11]. Humanized gnotobiotic animals can be used to test causative relationships between human microbiomes and host physiology while maintaining control over host genetics, diet and environment [12-16]. Components of IBD-associated microbiotas can enhance colitis severity in gnotobiotic mice [17-20], and complete microbiotas from donors both with and without IBD can induce intestinal inflammation in susceptible mice [19, 21-24]. However, the scale of these studies has precluded identifying specific functional consequences to the host of colonization with IBD microbiotas compared to healthy microbiotas. Here we use 30 human gut microbiotas from two independent cohorts recruited near New York, NY or near Los Angeles, CA, to demonstrate that IBD microbiotas drive distinct immune profiles in unchallenged gnotobiotic mice and more severe disease in colitis-susceptible gnotobiotic mice.

Since IBD pathophysiology is associated with a dysregulated T cell response [25], we asked whether IBD microbiotas could specifically alter CD4 T cell responses compared to healthy microbiotas. Thus we colonized germ free C57Bl/6J (B6) mice with fecal slurries or arrayed cultured fecal microbiota collections [26, 27] from two independent cohorts of healthy donors (n=11) or donors with IBD (n=13; Table S1) [8]. Four to six weeks post colonization, we performed a comprehensive measurement of CD4^+^ T helper (Th) and T regulatory (Treg) cells in the intestinal lamina propria of each colonized mouse using flow cytometry. While we found no difference in the proportion of IFNγ^+^ Th1, GATA3^+^ Th2, IL- 10^+^ or IL-17A^+^ CD4 T cells (Fig.1, S1), the average proportion of RORγt^+^Th (CD4^+^Foxp3^-^RORγt^+^) induced by IBD microbiotas was significantly higher than healthy donor-derived microbiotas in both colon and ileum (p=0.009, p=0.034 respectively; t-test; Fig. 1C, S2). RORγt^+^Th varied over an 6-fold range and included human microbiotas inducing similar proportions to common low/high Th17 reference communities (specific pathogen free (SPF) mouse microbiotas +/- segmented filamentous bacteria (SFB)) [4] (Fig.1D). The proportion of RORγt^+^Th was correlated with the proportion of IL-17A^+^ CD4 T-cells within each tissue (colon; p=2.3×10^−6^, R^2^=0.68, ileum; p=0.0018, R^2^=0.36; Fig. S1G).

**Figure 1:**
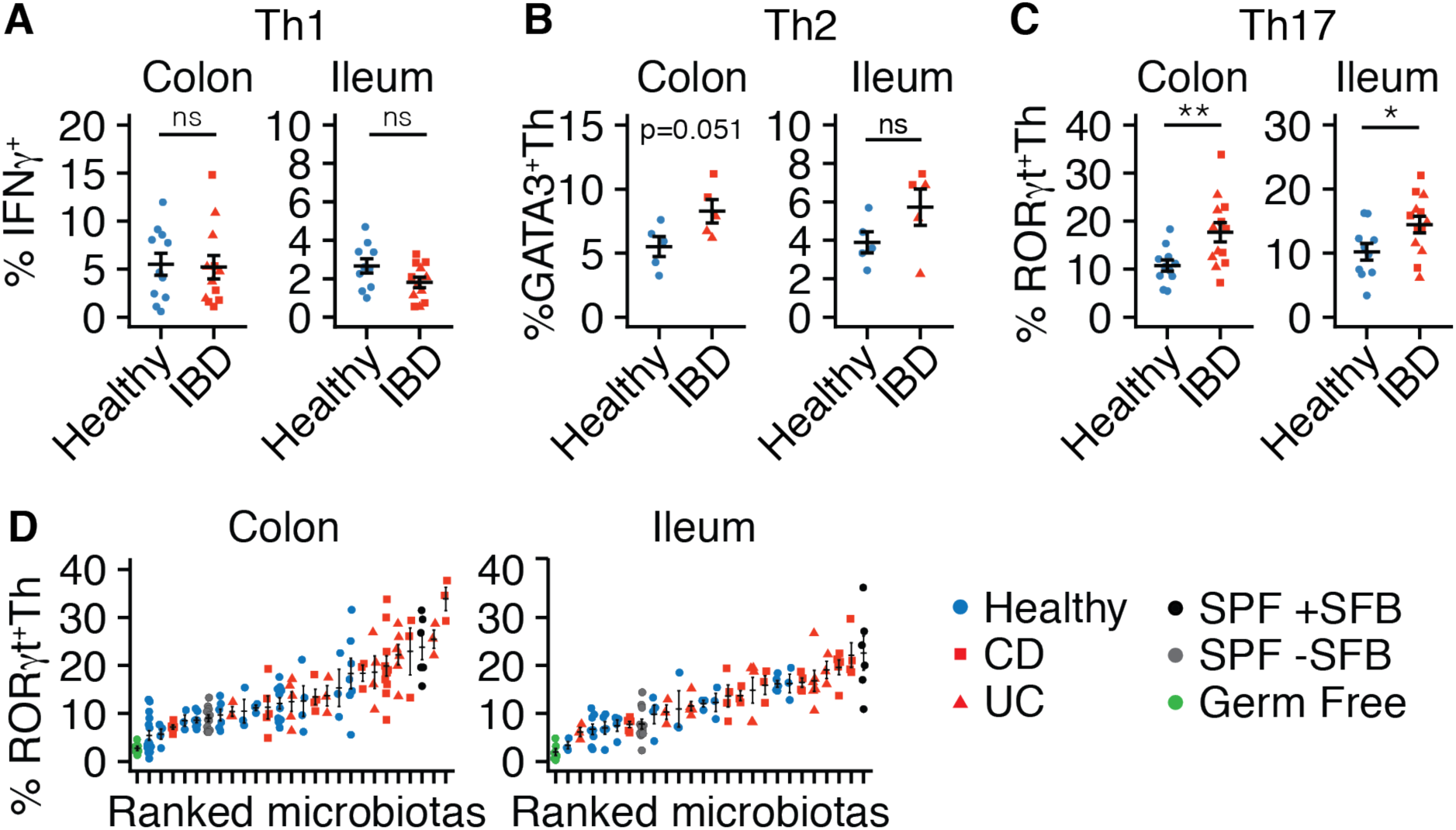
IBD-associated microbiotas enhance gut ROR γ t^+^ Th induction compared to healthy-associated microbiotas. The mean proportion of (**A**) IFNγ^+^, (**B**) GATA3^+^ and (**C**) RORγt^+^Th in colon and ileum of gnotobiotic mice colonized with IBD or healthy-donor microbiotas. Each symbol represents the mean value of a group of 3-12 mice colonized with a single microbiota; horizontal lines indicate mean ± SEM. n=11 healthy, 6 UC and 7 CD microbiotas (IFNγ and RORγt) n=5 healthy, 3 UC and 2 CD (GATA3); *p<0.05, **p<0.01. **D**. The proportion of RORγt^+^Th in colon and ileum varies across gnotobiotic mice colonized with fecal microbiotas from different human donors. Each symbol represents data from one mouse; horizontal lines indicate mean ± SEM. n=11 healthy, 6 UC and 7 CD microbiotas; *p<0.05, **p<0.01.

We hypothesized that the specific expansion of RORγt^+^Th may be the result of impaired FoxP3^+^Treg induction by IBD-associated microbiotas. As in previous studies [26, 28], the majority of microbiotas led to an expansion of gut FoxP3^+^Treg above baseline germ-free levels (Fig. 2A). Although variation in the proportion of FoxP3^+^Treg was significantly influenced by donor microbiota (p=1.8×10^−10^ (colon), p=4.9×10^−4^ (ileum); ANOVA), we observed no difference between the proportion of FoxP3^+^Treg induced by healthy or IBD- associated microbiotas in colon or ileum (Fig. 2B). We also found no correlation between the proportion of RORγt^+^Th and FoxP3^+^Treg induced by each microbiota in either tissue (Fig. S3). Recently, the gut-specific subset of Treg that co-express FoxP3 and RORγt (RORγt^+^Treg) was found to be microbiota-dependent and colitoprotective [29-33]. Induction of RORγt^+^Treg varied significantly with different microbiotas (p<1×10^−5^,ANOVA; Fig.2C-E). In contrast to the total Treg population, we observed a significant expansion of RORγt^+^Treg induced by healthy relative to IBD microbiotas in both colon and ileum (p<0.001, t-test; Fig. 2D-E). This difference was significant across both cultured and stool microbiotas (Fig.S4A) and across the two independent cohorts of IBD patients (Fig. S4B). Of note, the proportion of total FoxP3^+^Treg was only weakly correlated with RORγt^+^Treg in ileum (p=0.012; R^2^=0.24) and uncorrelated in colon (Fig. 2F), suggestive of a unique developmental pathway for RORγt^+^Treg. Helios^+^Treg were not differentially induced by healthy and IBD microbiotas and we observed only a weak correlation between RORγt^+^Treg and Helios^+^Treg (Fig. S5). It has been suggested that RORγt^+^Treg are uniquely positioned to regulate Th2 responses [32]. Although we observed a non-significant expansion of Th2 (GATA3^+^ FoxP3^-^ CD4^+^) cells in the colon of gnotobiotic mice colonized with IBD microbiotas relative to healthy microbiotas (p=0.051, t-test; Fig.1B), induction of Th2 cells was uncorrelated with RORγt^+^Treg (p=0.062, Fig.S5A). We also found no correlation between the proportion of RORγt^+^Treg and RORγt^+^Th or IFNγ^+^ CD4 T-cells (Fig. S6B-C).

**Figure 2:**
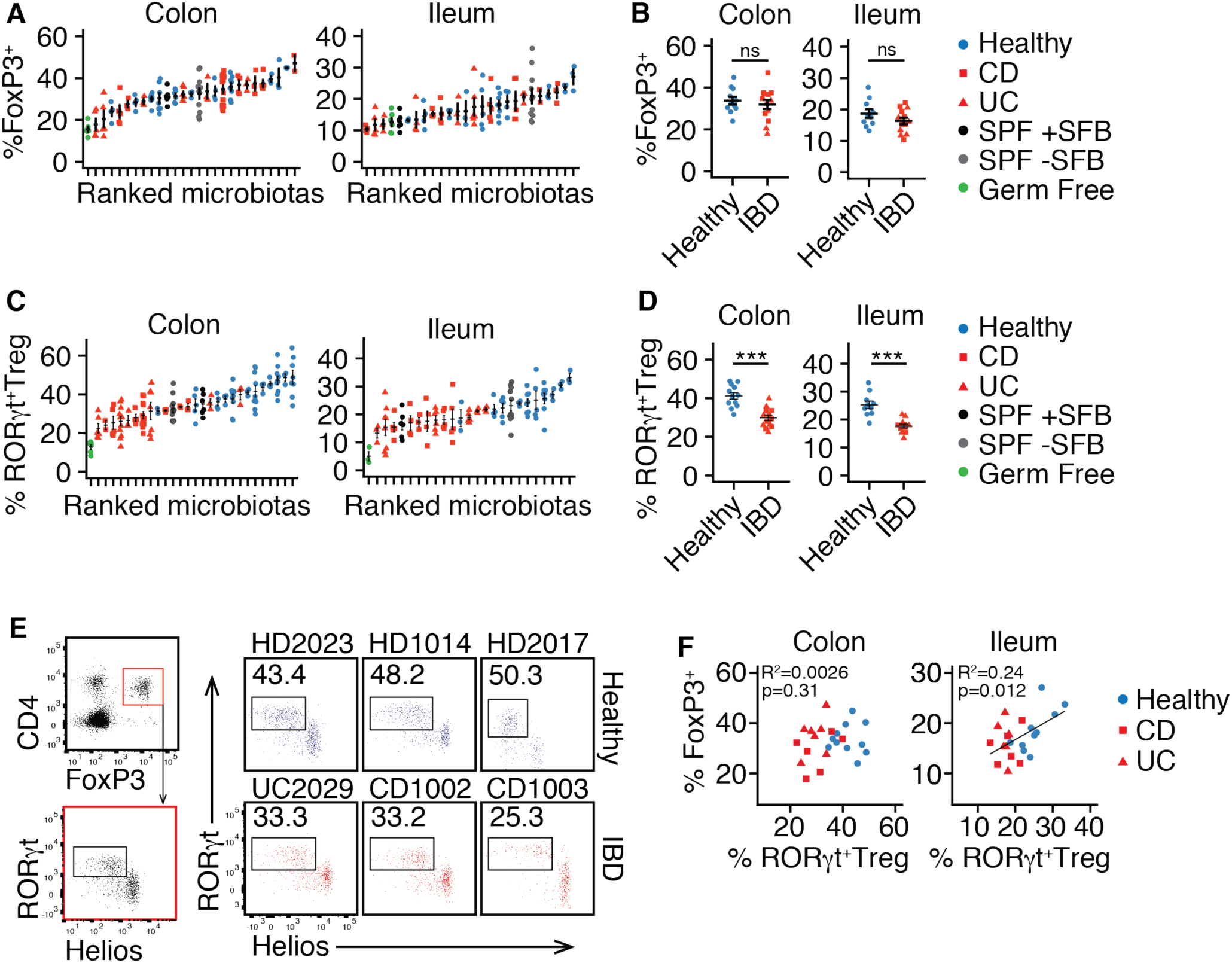
Healthy donor-associated microbiotas enhance ROR γ t^+^Treg compared to IBD-associated microbiotas. **A.** The proportion of FoxP3^+^Treg in the colon and ileum of gnotobiotic mice colonized with fecal microbiotas from multiple human donors. Each symbol represents data from one mouse; horizontal lines indicate mean ± SEM. **B.** The mean proportion of FoxP3^+^Treg induced by a donor microbiota is not significantly associated with the disease phenotype of the donor (ns – not significant; t-test). Each symbol represents the mean value from a group of 3-12 mice colonized with a single microbiota; horizontal lines indicate mean ± SEM. **C**. The proportion of RORγt^+^Treg in the colon and ileum of gnotobiotic mice colonized with fecal microbiotas from different human donors. Each symbol represents data from one mouse; horizontal lines indicate mean ± SEM. **D**. The mean proportion of RORγt^+^Treg in colon and ileum is increased in gnotobiotic mice colonized with healthy-donor microbiotas compared to IBD-donor microbiotas. Each symbol represents the mean value from a group of 3-12 mice colonized with a single microbiota; horizontal lines indicate mean ± SEM. n=11 healthy, 6 UC and 7 CD microbiotas ***p<0.001. **E**. Gating of RORγt^+^Treg and representative plots of colon RORγt^+^Treg in gntotobiotic mice colonized with six different microbiotas. **F**. Total FoxP3^+^ and RORγt^+^Treg are not significantly correlated in the colon and weakly correlated in the ileum. Each symbol represents the mean value from a group of 3-12 mice colonized with a single microbiota.

To assess if IBD-associated microbiotas influence colitogenesis, we tested healthy- and IBD-donor microbiotas in a gnotobiotic mouse model of colitis. Given the known importance of T-cells in IBD pathophysiology we chose a model of colitis that is both T cell- and microbiota-dependent. Transfer of CD45RB^HI^ (naïve) CD4 T-cells to Rag-deficient mice induces colitis-like pathology, but only in the presence of an immunogenic microbiota (hereafter the Rag T-cell transfer, (RagTCT) model) [34, 35]. 4-8 weeks prior to T-cell transfer, we colonized germ-free Rag1^-/-^ mice with fecal microbiotas from both healthy (n=16) or IBD human donors (n=14; see Data S1). As measured by loss in body mass, histology, and elevation of fecal lipocalin2 (LCN2), colitis was more severe in mice colonized with fecal microbiotas from individuals with IBD than those colonized with microbiotas from healthy donors (p=4.2×10^−5^, p=0.0058 at day 42 respectively for body mass and LCN2, t-test; Fig. 3). Loss in body mass was correlated with elevated fecal LCN2 (R^2^=0.33, p=1.4×10^−7^; Fig. S7). Remarkably, a significant difference in weight loss between healthy and IBD microbiotas was already detectable 7 days after T-cell transfer and became more prominent over time (Fig. S8). There was no significant difference in colitis severity between mice colonized with microbiotas from donors with UC compared to CD (p=0.59, t-test; Fig. 3D), and CD and UC microbiotas each independently induced colitis that was more severe than in mice colonized with healthy donor microbiotas (p<0.01, p<0.001 for UC and CD respectively, ANOVA; Fig. 3D). We replicated these findings in two independent cohorts of donors [8] (Fig. 3E, S9A). We also found both stool microbiotas and cultured collections of microbes from donors with IBD were similarly able to increase colitis susceptibility in mice, relative to healthy donor microbiotas (Fig. 3F, S9B). For ten donors, we assayed the colitogenicity of both the stool and cultured microbiota collection. Eight of the 10 cultured microbiotas transferred colitis of equivalent severity as the total stool microbiota derived from the same donor (Fig. S10).

**Figure 3:**
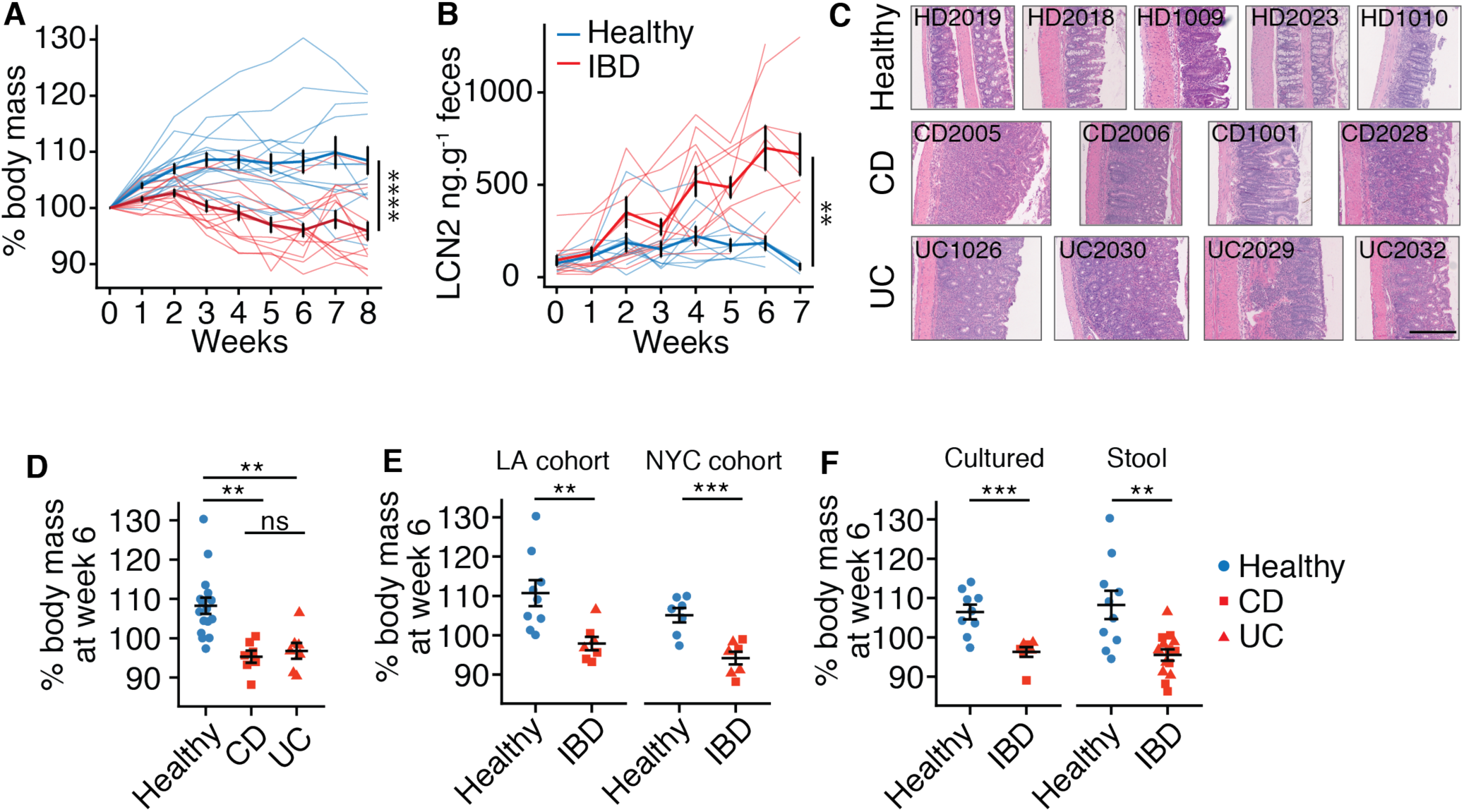
IBD-associated microbiotas transmit enhanced colitis susceptibility to mice. **A.** Body mass decreased and **B**. fecal lipocalin2 (LCN2) increased in RagTCT mice colonized with human microbiotas from healthy donors compared to IBD donors. Each thin line represents the mean data from a group of 5-15 mice colonized with a single microbiota (n=16 healthy donors assayed in a total 152 mice, n=12 IBD donors assayed in a total of 106 mice). Bold lines represent the mean ± SEM of all groups of mice colonized with either healthy donor or IBD donor microbiotas. **C**. Representative H&E stained colon sections from RagTCT mice colonized with different human donor microbiotas 5-7 weeks after T cell transfer. Scale bar = 200μm. **D**. We found no significant change in body mass loss between RagTCT mice colonized with human UC or CD microbiotas. Each point represents the mean weight change 6 weeks after T cell transfer of a group of 5-15 mice colonized with a single microbiota (n=6 CD donors, n=6 UC donors). Bold lines represent the mean ± SEM of all groups of mice colonized with either CD donor or UC donor microbiotas. **E.** IBD microbiotas from both donor cohorts induce more severe weight loss, as measured by weight loss at week 6, than healthy donor microbiotas. **F**. Cultured and stool IBD microbiotas both induce more severe weight loss, as measured by weight loss at week 6, than healthy donor microbiotas. **p<0.01,****p<0.0001

Finally, we sought to understand how the variation in CD4 T-cell responses we observed in unchallenged gnotobiotic B6 mice correlated with colitis severity in RagTCT mice colonized with the same donor microbiotas. A total of 9 healthy and 11 IBD microbiotas were tested in both models (Table S1). While colitis severity was not correlated with IFNγ^+^Th or FoxP3^+^Treg (Fig. 4A-B), the proportion of colon and ileum RORγt^+^Th induced by a microbiota in B6 mice was positively correlated with colitis severity in RagTCT mice colonized with the same microbiota (R^2^=0.46, p=9×10^−4^; R^2^=0.34, p=0.006 for colon and ileum respectively; Fig. 4C). Induction of RORγt^+^Treg in B6 mice was inversely correlated with colitis severity in RagTCT mice (R^2^=0.47, p=0.0007; R^2^=0.43, p=0.002 for colon and ileum respectively; Fig. 4D). The proportion of IL-17A^+^ CD4 T-cells induced in colon was also weakly associated with colitis severity (R^2^=0.26, p=0.015; Fig. S10A). Colitis severity was not associated with Helios^+^Treg, IL-10^+^, or GATA3^+^ CD4 T-cells (Fig. S11). Hierarchical clustering of microbiotas based upon only the proportion of RORγt^+^Th and RORγt^+^Treg in the colon and ileum of unchallenged B6 mice was sufficient to correctly predict the health status of 20/22 (90.9%) donors (Fig. 4E). Induction of these cell populations in B6 mice was also predictive of colitis severity in RagTCT mice colonized with the same microbiota, independent of microbiota donor health (Fig. 4F). A linear model explained 70% of the variation in colitis severity (weight loss at week 6) as a function of the proportion of both RORγt^+^Th and RORγt^+^Treg in colon (R^2^=0.70, p=2.1×10^-5^; F-test).

**Figure 4:**
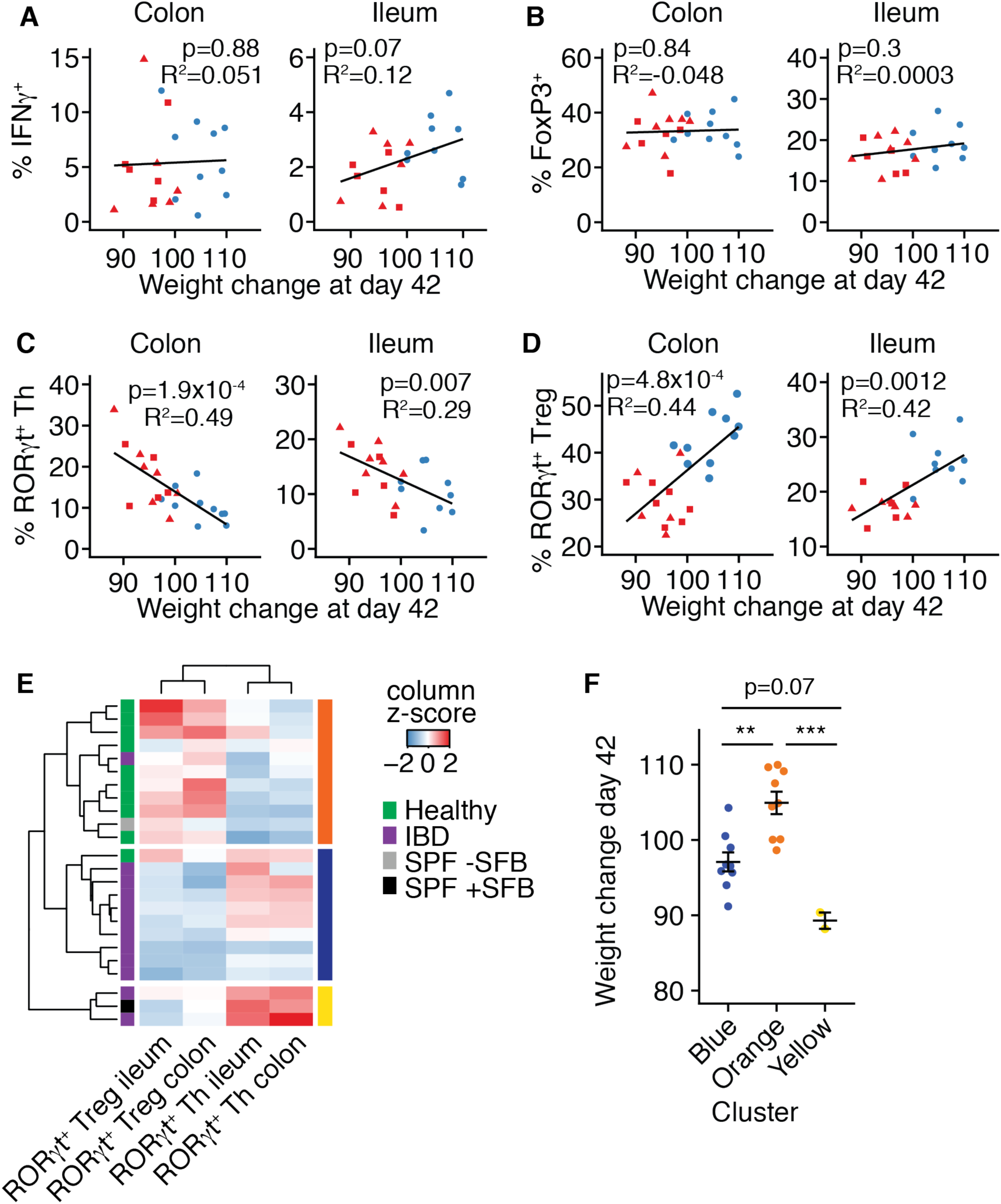
Homeostatic induction of ROR γ t^+^Treg and ROR γ t^+^Th predicts experimental colitis severity and human microbiota donor health. **A.** IFNγ^+^ Th and **B**. FoxP3^+^Treg in the colon and ileum of B6 gnotobiotic mice are not correlated with colitis severity (as measured by weight loss) of RagTCT mice colonized with the same microbiotas. Homeostatically induced **C.** RORγt^+^Th and **D.** RORγt^+^Treg in the colon and ileum of B6 gnotobiotic mice are correlated with colitis severity (as measured by weight loss) of RagTCT mice colonized with the same microbiotas. **E.** Hierarchical clustering of donor microbiotas based on the proportion of RORγt^+^Treg and RORγt^+^Th induced in colon and ileum of gnotobiotic B6 mice identifies three clusters which related to donor health. The left color bar indicates microbiota human donor health and the right color bar denotes the three clusters for microbiota tested in the RagTCT model. **F**. Clusters were also related to colitis severity (as measured by weight loss) in RagTCT mice colonized with the same microbiotas. The color of each point indicates its cluster in pane C. The body weight data represents the mean measurements of groups of 5-15 mice colonized with a single human donor microbiota ± SEM. p values are calculated by ANOVA with Tukey‘s correction for multiple comparisons. **p<0.01, ***p<0.001

Using two donor cohorts we show that human microbiotas transferred to gnotobiotic mice alter homeostatic immune responses and the severity of gut pathology in susceptible mice in ways that are highly predictive of the original human host disease. This suggests that, in addition to host genetics, the composition of the gut microbiota is a significant risk factor for complex disease. Furthermore, it provides an etiological hypothesis for a role of the gut microbiota in IBD risk whereby individuals harboring communities that enrich tolerogenic RORγt^+^Tregs are a lower risk, while those harboring communities that enrich RORγt^+^Th are at increased risk. Microbiota analysis by 16S rDNA amplicon sequencing did not distinguish between the fecal microbiotas of healthy donors and donors with CD or UC (p=0.58, PERMANOVA; Fig. S12) suggesting the functional impact of the microbiota lies in the strain-level composition of each unique microbiota. Cultured collections of microbes from donors with IBD recapitulate observations in mice colonized with complete fecal microbiotas, providing the prospect of determining the relative contribution of individual isolated strains to specific host phenotypes [26]. Finally, these data further support the microbiota as a viable target for therapeutic intervention in IBD, and provide a hypothesis for rational design of microbiota-directed preventative strategies and treatment in IBD.

## Materials and Methods

Unless otherwise stated, chemicals were obtained from Sigma Aldrich.

### Human samples and bacterial culture

Human stool samples from were frozen at −80°C before processing. Samples were pulverized under liquid nitrogen. Under strict anaerobic conditions ∼∼500mg of pulverized stool from each donor was blended into a slurry (40-50mg/ml) in pre-reduced bacterial culture media (LYBHIv4 media [36]; 37g/l Brain Heart Infusion [BD], 5g/l yeast extract [BD], 1g/l each of D-xylose, D-fructose, D-glactose, cellubiose, maltose, sucrose, 0.5g/l N-acetylglucosamine, 0.5g/l L-arabinose, 0.5g/l L-cysteine, 1g/l malic acid, 2g/l sodium sulfate, 0.05% Tween 80, 20μg/ml menadione, 5mg/l hemin (as histidine-hemitin), 0.1M MOPS, pH 7.2). The slurries were passed through sterile 100μm strainers to remove large debris. To store for later administration to mice, slurries were diluted 1:20 in LYBHIv4 media containing 15% glycerol (final concentration) and stored at −80°C.

Arrayed culture collections were generated for selected donors as previously described. Briefly, clarified and diluted donor stool was plated onto a variety of solid selective and non-selective media under anaerobic, micro-aerophilic and aerobic conditions. Plates were incubated for 48-72 hours at 37°C. 384 single colonies from each donor microbiota were individually picked and regrown in liquid LYBHIv4 media for 48 hours under anaerobic conditions. Regrown isolates were identified at the species level using a combination of MALDI-TOF mass spectrometry (Bruker Biotyper) and 16S rDNA amplicon sequencing. An average of 16 unique species were isolated from each fecal sample (range 10-29; Table S2). There was no significant difference in the number of unique isolates obtained from healthy donor and IBD donors (p=0.19, Mann-Whitney). All regrown isolates were stored in LYBHIv4 with 15% glycerol at −80°C as both arrayed culture collections in 96 well plates and pooled as a cocktail for administration to mice.

### Gnotobiotic mice

Germ free C57Bl/6J and C57Bl/6J Rag1-deficient (B6.129S7-Rag1^tm^1^Mom^/J) mice were bred in-house at the Mount Sinai Immunology Institute Gnotobiotic Facility in flexible vinyl isolators. To facilitate high-throughput studies in gnotobiotic mice we utilized “out-of-the-isolator” gnotobiotic techniques [26]. Shortly after weaning (28-42 days old) and under strict aseptic conditions, germ-free mice were transferred to autoclaved filter-top cages outside the of the breeding isolator and colonized with human microbiotas; Mice were colonized with 200-300μl of a fecal slurry or pooled cocktail of cultured strains by oral gavage, given only once. All experiments were performed at least 28 days after colonization. All phenotyping experiments in B6 mice included both male and female mice. Of the 30 microbiotas screened in the Rag1-deficient colitis model, 26 were tested in both male and female mice. Animal procedures were approved by the Institutional Animal Care and Use Committee (IACUC) of Mount Sinai School of Medicine.

### 16S rDNA sequencing and analysis

The composition of human and mouse fecal samples was analyzed by 16S rDNA amplicon sequencing as previously described [37-39]. DNA was extracted by bead-beating followed by QiaQuick columns (Qiagen) and quantified by Qubit assay (Life Tehcnologies). The V4 region of the 16S gene was amplified by PCR and paired-end 250bp reads sequenced on an Ilumina MiSeq. Analysis was performed with MacQIIME 1.9.1.8 and custom R scripts. OTUs were picked with 97% sequence similarity. OTUs were aligned to the Greengenes reference set, requiring 150bp minimum sequence length and 75% ID.

### Lymphocyte isolation

Spleen and mesenteric lymph nodes were collected into RPMI containing 5% fetal bovine serum (FBS). Single cell suspensions were obtained by pressing though 40μm strainers.

Red blood cells were removed with ACK Lysing Buffer (Gibco). Gut tissues were separated, opened longitudinally and washed in Hanks Buffered salt solution (HBSS) to remove intestinal contents. Ileum was defined as the distal 1/3 of the small intestine. Peyers patches were removed. Epithelial cells were removed by gentle shaking in HBSS (Ca/Mg free) with 5mM EDTA, 15mM HEPEs and 5% FBS for 30 minutes. The remaining tissue was washed in HBSS before mincing with scissors into digestion buffer (HBSS, 2% FBS, 0.4mg/ml Collagenase Type IV [Sigma Aldrich C5138], 0.1-0.25mg/ml DNase1 [Sigma Aldrich DN25]) and incubated at 37°C with gentle shaking for 30-40 minutes. The resulting suspensions were passed sequentially though 100μm and 40μm strainers. No gradient centrifugation enrichment of lymphocytes was performed.

### Flow cytometry

For analysis of intracellular cytokines (IL-10, IFNγ, IL-17A), lamina propria lymphocytes were restimulated in complete RPMI with 5ng/ml phorbal 12-myristate 13-acetate (PMA) and 500ng/ml ionomycin in the presence of monensin (Biolegend) for 3.5 hours at 37°C. Dead cells were excluded from all analyses using Zombie Aqua Fixable Viability dye (Biolegend). For intracellular cytokine staining, cells were fixed with IC Fixation Buffer (eBioscience) and transcription factors were detected in unstimulated cells fixed with FoxP3 Fixation/Permeabilization buffers (eBioscience). Antibodies used in this study are listed in Table S2. All data was acquired on the same LSRII instrument (BD Biosceinces) and analyzed using FloJoX (TreeStar).

### T cell transfer colitis

T cell transfer colitis experiments were performed as previously described[40]. Briefly, naïve (CD45RB^HI^, CD25^-^) CD4 T cells were isolated from the spleen and subcutaneous lymphnodes of 7-9 week old specific pathogen free C57Bl/6J mice (The Jackson Laboratory). Following tissue dissociation and red blood cell lysis CD4^+^ T cells were enriched using negative magnetic selection (Magnisort, eBioscience). The resulting cells were stained for expression of CD4, CD25 and CD45RB. A fraction representing ∼50% of the total CD4^+^ population, selected on the basis of absent CD25 staining and high CD45RB staining was sorted using a FACSAria (BD Biosciences). Purity of the sorted fraction was checked and routinely exceeded 98%. Sorted cells were washed multiple times with sterile PBS. Gnotobiotic Rag1^-/-^ mice received 1×10^6^ CD45RB^HI^ T cells in 200μl of sterile PBS by intraperitoneal injection. Donor cells were sex-matched to recipients.

Mice were weighed and fecal pellets were collected at the time of T cell transfer and weekly thereafter. Any mouse experiencing >80% loss in body weight or which was deemed otherwise moribund was euthanized. In these cases, the last measurements of body mass or LCN2 taken for that mouse were carried forward and included in the data for subsequent time points. Inter-experimental variation was assessed across the screen by including one cultured donor microbiota in each experiment. This donor (UC1024) induced highly reproducible colitis in many repeats over ∼2 years (Fig. S12). We set pre-determined exclusion criteria for any experiment where mice colonized with UC1024 did not develop colitis in this reproducible manner.

### LCN2 measurements

Lipocalin2 concentrations were measured in feces as a biomarker of intestinal inflammation [41]. Fecal pellets were collected into sterile pre-weighed and barcoded tubes and frozen at −20°C until the time of analysis. Pellets were weighed and suspended in 500μl of sterile PBS by shaking in a BeadBeater (with no beads in the tube) for 2 minutes. Tubes were centrifuged at 4000rpm for 20 minutes. The resulting supernatant was assayed for LCN2 by sandwich ELISA (R&D systems). The concentration of LCN2 was normalized to the weight of the input feces.

## Acknowledgements

We thank C. Fermin, E. Vazquez and G. Escano of the Mount Sinai Immunology Institute Gnotobiotic Facility for exceptional technical support.

## Author contributions

G.J.B. and J.J.F. conceptualized the project. G.J.B., E.J.C., O.H.V., S.R.L., R.N. and A.M. performed experiments and analyzed the data. I.M. and Z.L. generated cultured collections of microbes. M.M., A.D., D.G., D.P.B.M., N.S., J.B., J.P.J., J.C.C., A.G., B.E.S., J-F.C. and M.C.D. provided essential research resources. G.J.B and J.J.F wrote the manuscript with input from all authors. J.J.F. supervised the study.

## Data availability

16S rDNA datasets analyzed in the manuscript are available through NCBI under accession number PRJNA403997.

